# Alphafold2 refinement improves designability of large de novo proteins

**DOI:** 10.1101/2024.11.21.624687

**Authors:** Christopher Frank, Dominik Schiwietz, Lara Fuß, Sergey Ovchinnikov, Hendrik Dietz

## Abstract

Recent advances in computational protein design have enabled the creation of novel proteins for a variety of purposes. The capability for producing custom-shape high-quality backbones for very large proteins would further expand the scope of protein design. To this end, here we introduce the AlphaFold2 (AF2) cycler (af2cycler) design pipeline. AF2cycler is used to refine draft protein backbone geometries of arbitrary shape created by the Chroma diffusion model or other methods, using a combination of AF2 and ProteinMPNN to achieve high designability with minimal deviations from the initial target structure. In silico testing on multiple protein designs (100-1000 amino acids) demonstrated improvements in structural integrity and designability after af2cycling confirmed by enhanced ESMFold repredictions. Experimental wet lab validation through the design of a variety of test proteins with distinct shapes, each comprising 1000 amino acids, showed structural agreement between in silico predictions and transmission electron microscopy (TEM) imaging, establishing the af2cycler’s efficacy in translating designs into real-world results. Af2cycler provides a convenient and reliable protein design workflow, particularly for large proteins, with potential for expanding to applications in areas such as design of higher-order protein complexes and multi-state backbone optimization.

## Introduction

*De novo* protein design, which aims to create entirely new proteins from first principles, has become an important focus in computational biology (*1, 2*). This field promises transformative applications in biotechnology, medicine, and synthetic biology (*1, 2*). The recent awarding of the Nobel Prize for AlphaFold2 highlighted its groundbreaking ability to predict protein structures with atomic accuracy, reshaping our understanding of protein folding. While originally developed for structure prediction, AlphaFold2 (*3*) has also demonstrated its utility in *de novo* protein design (*4–8*), especially in generating binders (*7*) and high-quality backbones, particularly for large proteins (*4*). Despite these advancements, existing approaches to *de novo* protein design, including diffusion and flow matching methods (*9*–*15*), often struggle when tasked with designing larger proteins. Although notable exceptions, such as Proteus (*16*), have shown success in this space, current approaches may yield designs with suboptimal backbone configurations, including over-compactness and irregular geometries.

Chroma (*9*), a recently developed computational approach for de novo protein design, offers a flexible and intuitive platform for generating complex protein structures. Chroma’s programmability allows users a convenient approach for specifying desired structural features, such as shape or secondary structure content, while maintaining the freedom to explore a vast conformational space. Its user-friendly interface and broad adaptability make it accessible to a wide range of researchers, enabling rapid prototyping of custom proteins with minimal computational expertise. Although Chroma generates visually promising large protein structures that avoid compactness issues that may arise in methods like RFDiffusion (*14*) (Supp. Fig. 1 A & B), further analysis using ESMFold (*17*) reveals that Chroma-generated designs tend to exhibit low designability when subjected to more rigorous structural criteria (TM Score > 0.85) (Supp. Fig. 1 C). Subsequent solubleMPNN-based (*18*) sequence redesign does not increase the in-silico success rate significantly (Supp. Fig. 1 C).

We hypothesized that optimizing backbone quality of Chroma-generate protein design drafts would help addressing these issues and translate into reduced risk of experimental failure. In previous work AF2 showed great potential for the design of large proteins with high experimental success when combined with ProteinMPNN (Supp. Fig. 1 D). We thus developed a pipeline termed the **af2cycler** that leverages AlphaFold2’s intrinsic denoising capabilities in combination with ProteinMPNN to refine Chroma-generated backbones, leading to significantly enhanced designability with minimal alterations to the original design target, which we refer to as the “initial draft backbone”.

The protein design pipeline consists of three key steps: First, initial protein design drafts are generated using the Chroma diffusion model, as previously described (*9*). Chroma can utilize a variety of conditions like shapes, symmetries or sequences and generate custom protein structures. Chroma Design is then used to generate a sequence. Both the sequence and the corresponding structure are then processed through the af2cycler, the central piece of this pipeline (Fig. 1). In this stage, the Chroma-generated backbone undergoes a refinement process using AlphaFold2 and ProteinMPNN, designed to improve its structural integrity and designability (Fig. 1 green box). The final step involves first sequence generation using solubleMPNN (*18*) and subsequent screening of the sequences with ESMFold to assess their folding accuracy and selecting the most promising candidates for experimental validation.

**Figure 1:**
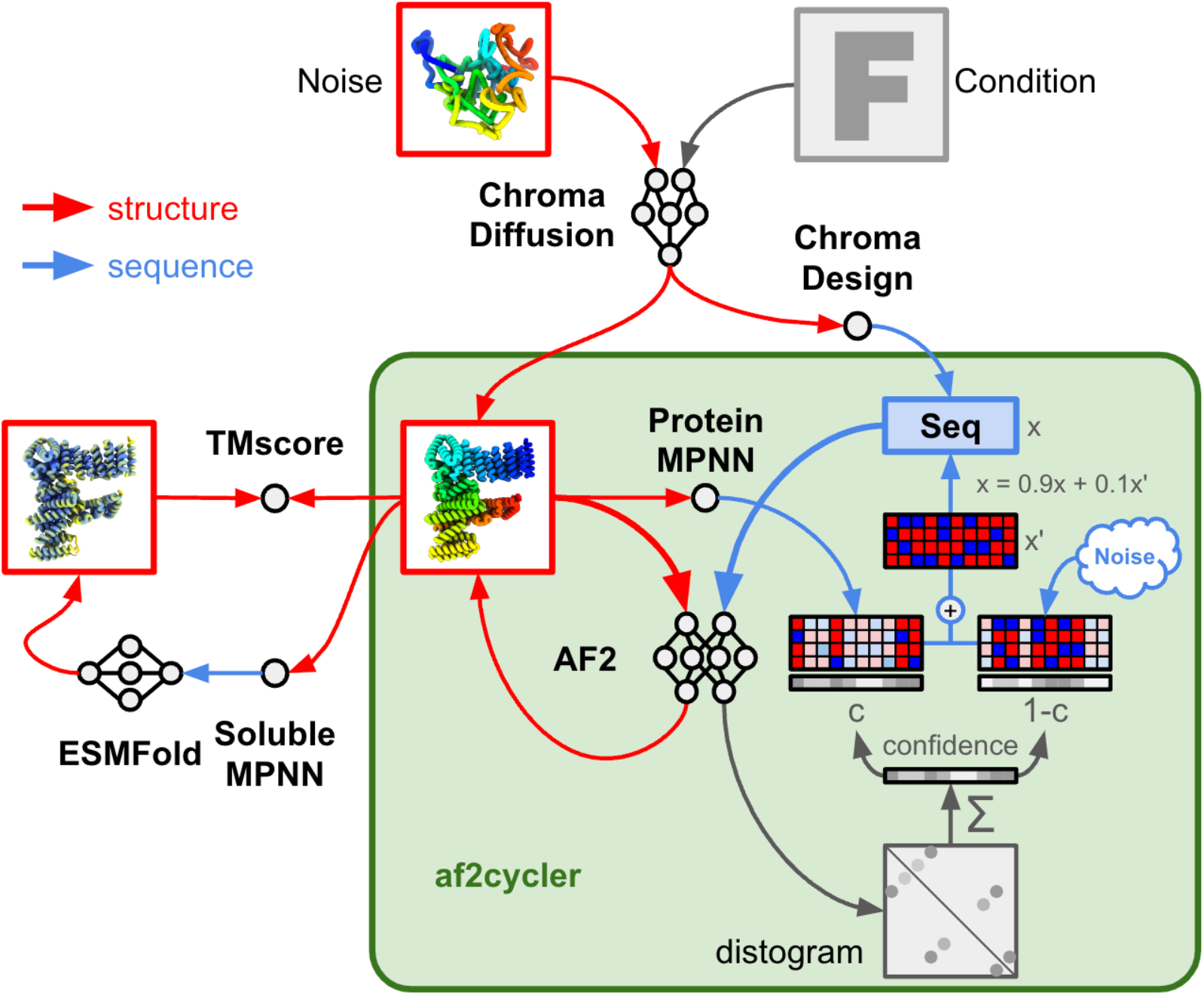
The af2cycler: Schematic representation of the af2cycler. Red arrows indicate the flow of structural information, while blue arrows indicate the flow of sequences. The af2cycler works by first using Chroma to design proteins that satisfy a certain condition. Secondly, the resulting structures as well as the designed sequences (with ChromaDesign) are put into the af2cycler main loop. The sequence is predicted with AF2 using the structure as a template. The resulting structure is used for the next round, as well as an input to ProteinMPNN. The distogram from the AF2 prediction is used to calculate a per residue confidence, capturing the number of contacts each residue has. The ProteinMPNN logits are then scaled with this confidence, while the compliment (1-C) is used to scale gumbel noise. The sum of these two is added to the one hot sequence and another argmax operation is performed, before putting this sequence back into AF2. Finally, the resulting structure is subject to solubleMPNN sequence design and a final ESMFold prediction.

The af2cycler, illustrated in Fig. 1, leverages AF2’s intrinsic ability to start with a template as an initial guess and locally descend the learned energy potential to refine input structure (*19*). In this process, the Chroma-designed protein “draft” sequence, without coevolutionary data, is fed into AF2. The draft structure generated by Chroma is provided as a template, initial guess (*20*) and used to initialize the structure module (*21*). One pass through the AF2 network is performed (zero-recycle prediction) and because a template matching the sequence is provided, this initial AF2 prediction is only slightly changed when compared to the original backbone, correcting structural inaccuracies without major alterations. This step can be understood as a single “denoising step.”.

Following this, ProteinMPNN is employed to generate a new sequence based on the refined structure (AF2 output). The ProteinMPNN sequence logits are then scaled by introducing a per-residue confidence (C) that is derived from the sum of contacts from the AF2 distogram. This scaled MPNN logits are summed with gumbel noise scaled by the complement of the confidence (1-C). Finally, the original sequence is updated by adding this noise augmented MPNN sequence and an argmax operation is performed before the sequence is input again into AF2. The confidence weighting is designed to assess how well-packed each residue is, adding more noise to under-packed residues and amplifying the weight of the ProteinMPNN logits in residues with sufficient contacts. This approach is based on the hypothesis that high-quality backbones will produce well-defined ProteinMPNN sequences, which converge toward a consensus sequence that encodes the backbone structure effectively.

The refined sequence, along with the updated AF2 prediction, is then cycled back into AF2 for additional refinement. This cycle of AF2 prediction and ProteinMPNN redesign is repeated for 10 to 20 iterations, or until the design reaches convergence, as determined by confidence metrics such as pLDDT and PAE.

The refined structure generated by the af2cycler can then undergo a final redesign using solubleMPNN to produce a completely new sequence. This sequence is subsequently validated by ESMFold, which repredicts the folding of the sequence into the intended structure. By integrating this final validation step, we ensure that the newly designed protein will adopt the desired conformation. Af2cycler harnesses the high programmability and user-friendly nature of Chroma, creating an effective, versatile, and reliable pipeline for the design of large proteins. Af2cycler enhances the draft designs by correcting structural inaccuracies, making this approach particularly suitable for designing large, complex proteins that maintain their stability and functionality.

### Computational benchmarking

To evaluate the impact of the af2cycler on protein design quality, we generated a test set of 50 proteins per length ranging from 100 to 1000 amino acids (AA) using Chroma (1000 steps, no additional conditioners). The af2cycler was then applied to these designs, performing 10 denoising steps on each. Notably, the af2cycler made only slight modifications to the backbone structures (Fig. 2A), with minimal deviations for smaller proteins, while larger designs exhibited a greater increase in root mean square deviation (RMSD) and TM Score (Fig. 2 B & Supp. Fig. 2 A). This increase primarily resulted from larger rearrangements in elongated domains (Fig. 2A). Despite these changes, the overall TM-score remained high (Supp. Fig. 2 A), even for designs up to 1000 AA, indicating strong structural similarity to the original Chroma designs.

**Figure 2:**
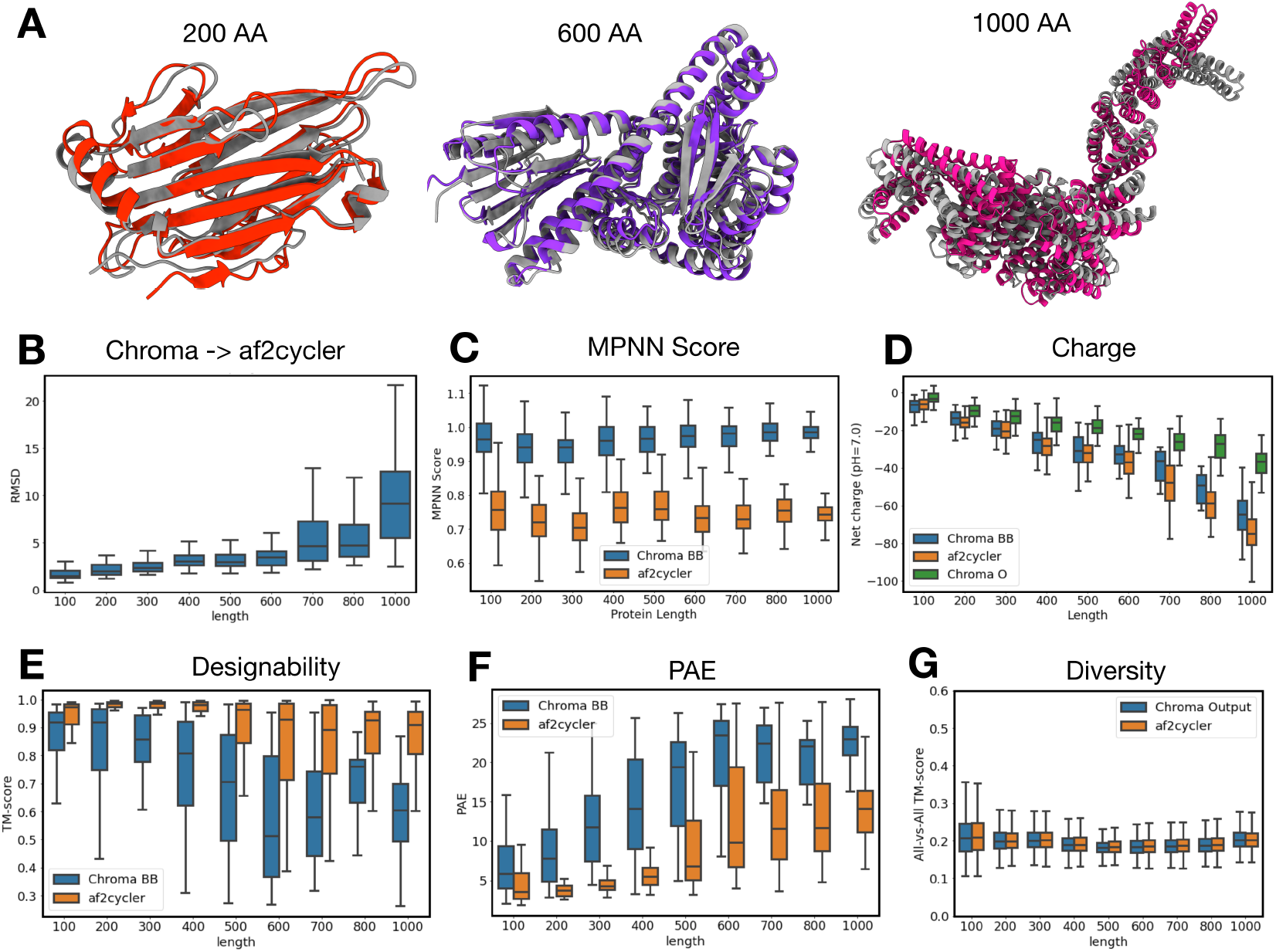
In silico benchmarking: A) Overlays of Chroma designed structures (grey) to af2cycler output (color) for different amino acid (AA) lengths. B) Boxplot showing the RMSD between the Chroma designed backbones and the af2cycler output for different lengths. C) Boxplot showing the MPNN score for Chroma or af2cycler treated backbones. D) Total net charge at pH=7.0 for sequences designed with solubleMPNN on Chroma or af2cycler treated backbones. Chroma Original sequences are shown in green as a reference. E) Designability (TM-score) of soluble MPNN designed sequences for Chroma designed or af2cycler treated backbones predicted with ESMFold. F) Predicted Alignment Error (PAE) for the predictions made in E) with ESMFold. G) Generated diversity (All-vs-All TM-score) of Chroma designed and af2cycler treated backbones

Following the af2cycler treatment, we generated sequences for both the original Chroma backbones and the af2cycler-refined backbones using solubleMPNN. We observed a significant reduction in MPNN scores for the af2cycler-treated designs compared to those generated directly from the Chroma backbones (Fig. 2 C), indicating that the cycled backbones represented higher-quality, more idealized structures. Additionally, we calculated the net charge of the designed sequences and found that the af2cycler-treated backbones had a lower net charge compared to the untreated designs (Fig. 2 D). Interestingly, both groups of solubleMPNN-designed sequences were more negatively charged than those generated by Chroma alone, likely due to the low sampling temperature (0.1) used in solubleMPNN.

We further evaluated the designability of these sequences by repredicting their structures with ESMFold. The af2cycler-treated backbones demonstrated significantly higher designability, with median TM-scores exceeding 0.85, even for the largest 1000 AA designs (Fig. 2E) and a median RMSD of 5 Å (Supp. Fig. 2 B). These predictions also showed lower predicted alignment error (PAE) (Fig. 2F) and higher pLDDT (predicted local distance difference test) (Supp. Fig. 2 C), further supporting the high success rate for large designs. Importantly, analysis of the designed structures revealed that the af2cycler did not reduce the overall complexity of the proteins (Fig. 2G), as the all-vs-all TM-scores of the designs were unaffected by the denoising process. Surprisingly, despite the strong bias towards helical structures in larger designs (Supp. Fig. 2, D), the overall structural diversity remained high even at larger sizes (Fig. 2G). These findings demonstrate that the in-silico metrics of Chroma-generated structures can indeed be enhanced by treatment with the af2cycler.

### Experimental investigations

For laboratory testing, we chose to af2cycler-design proteins consisting of 1000 amino acids, with tertiary structures approximating different letters of the Latin alphabet, as this “shape conditioning” feature was readily available in the Chroma software package. We expected these distinctive shapes to be easily identifiable in negative stain transmission electron microscopy (nsTEM), aiding structural investigations. For each letter of the Latin alphabet, we designed a set of five draft designs using Chroma with shape conditioning. These drafts were then subjected to 10 iterations of the af2cycler, followed by sequence generation with solubleMPNN and structural validation using ESMFold. As above, the af2cycler led to improvements of the backbone quality as manifested by reduced RMSD to design targets and reduced overall PAE of the predictions (Fig. 3 A).

**Figure 3:**
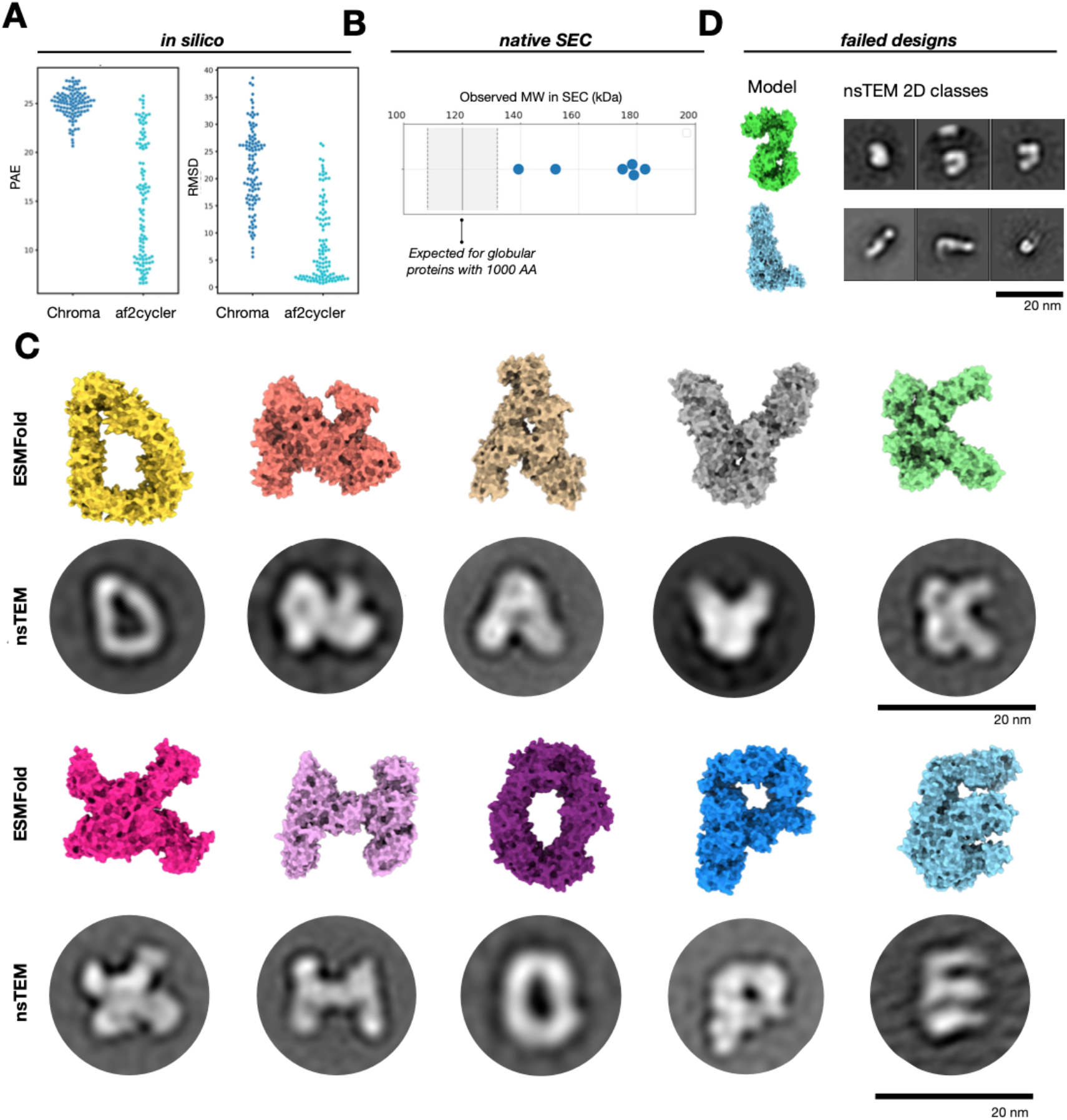
Experimental benchmarking of letter design: A) Swarmplot showing the reduction in RMSD and PAE for ESMFold predictions generated from sequences designed with Chroma of after af2cycler treatment. B) Observed MW in kDa on a calibrated native SEC for six letter shaped proteins with the expected MW region highlighted C) (Top Row) surface view renderings of the ESMFold predictions of the experimentally tested proteins. (Bottom Row): Best selected reference free 2D class average obtained with RELION of nsTEM data D) failed designs in nsTEM. Surface renderings are ESMFold models

We manually selected protein candidates considering high confidence scores. Proteins that did not maintain their intended shapes or showed poorly distinguishable simulated 3D volumes (such as ‘M’ or ‘B’) were excluded from testing, as we expected to experience difficulties in resolving their structures in negative-staining TEM imaging. Twelve proteins with shapes corresponding to the letters D, N, A, V, K, X, H, O, E, P, S and L were selected and expressed in E. coli and purified using Nickel-NTA chromatography.

All constructs expressed and SDS-PAGE gel electrophoresis confirmed the correct molecular weight (MW) of the proteins. The purity after Nickel-NTA chromatography purification was sufficient for further analysis. For the six proteins with the highest yield, we also performed native size exclusion chromatography (SEC) and found that the measured MWs were larger than expected, suggesting that the proteins adopted a non-globular shape that deviated from the standard radius assumed by the SEC calibration (Fig. 3 B). We imaged the proteins subsequently using nsTEM and found homogenous, monomeric protein distribution (Supp. Fig. 3). We computed 2D class averages from individual particle images using RELION (*22*). For 10 out of the 12 tested proteins, the resulting 2D classes showed strong agreement with the expectations from the designed proteins, successfully representing the intended letter shapes (Fig. 3 C and Supp. Fig. 4). While certain proteins, such as ‘H,’ exhibited some flexibility, as indicated by more diverse 2D classes, most proteins predominantly maintained their designed shapes. Only for the ‘S’ and ‘L’ shaped proteins the 2D classes from nsTEM analysis did not result in strong agreement to the designed shape (Fig. 3 D).

These findings demonstrate the viability of the af2cycler pipeline to rapidly prototype custom-shape large proteins with high likelihood of successful experimental outcome.

### Pushing the size limit of protein design

Encouraged by these initial results, we explored the size limitations of the Chroma-af2cycler design pipeline. We observed that the final confidence scores, specifically pLDDT and PAE, did not reach as high levels anymore for protein designs comprising more than 1000 amino acids (AAs) in length. This suggests that AlphaFold2 may be approaching its functional limits for accurately predicting structures at this scale, even when provided with a template, an initial guess, and all-atom initialization. However, despite this limitation, the af2cycler was still able to generate backbones that could be used for subsequent sequence design with ProteinMPNN even for proteins comprising up to 1500 AA, as exemplified with a triangular protein with 1500 AAs that we drafted in Chroma and subjected to 20 iterations of af2cycler refinement. AF2, supplemented with initial guess (*20*) and all atom initialization (*21*) struggled to accurately predict this large structure, yielding a root mean square deviation (RMSD) of 10 Å. However, AlphaFold3 (AF3-server) (*23*) was able to predict the structure with higher accuracy (Fig. 4 A). Due to constraints in the computational resources available to us, we were unable to run ESMFold on sequences longer than 1000 AAs. Experimentally, we tested a candidate sequence for the 1500-AA triangle design. The protein was expressed using mild conditions, and purification via Ni-NTA chromatography yielded a clean sample with the expected molecular weight (MW), as confirmed by SDS-PAGE (Fig. 4 B). Subsequent imaging with nsTEM revealed globular proteins, with triangle-like shapes. Reference-free 2D class averages generated from nsTEM data, showed good agreement with the design within the resolution, and an initial 3D reconstruction produced a 3D volume that overlayed well with the AF3 prediction (Fig. 4 B). These findings demonstrate that the Chroma-af2cycler pipeline has the potential to be extended to accommodate protein designs as large as 1500 AAs and possibly beyond.

**Figure 4:**
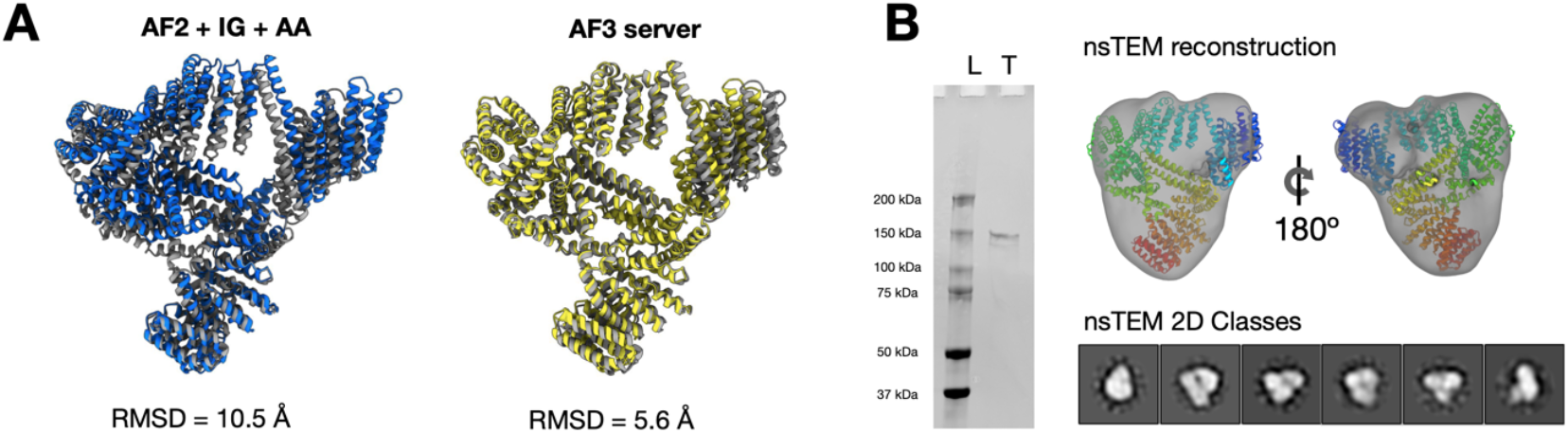
Experimental characterization of a 1500 AA protein. A) Af2cycler output (grey) overlaid with either AF2 with Initial guess (IG) and all atom initialization (AA) or AF3 (server) output of the best solubleMPNN designed sequence. B) Experimental investigation of 1500 AA protein: (left) SDS-PAGE gel showing Biorad precision dual color plus protein ladder (L) and the purified 1500 AA protein (T). (Right) exemplar nsTEM 2D classes and 3D reconstruction of the 1500 AA protein

## Discussion

De novo protein design has emerged as a frontier in computational biology, driven by the ambition to create new proteins that meet specific functional and structural criteria. Diffusion-based methods, like Chroma, offer an intuitive and flexible platform for designing complex proteins, but challenges remain in terms of improving the in silico and in vitro success. The af2cycler pipeline addresses these limitations by combining AF2’s denoising capabilities with the sequence design potential of ProteinMPNN. The af2cycler refines initial draft designs from Chroma or other input to improve their designability, yielding higher-quality structures suitable for further computational work and experimental validation. Our computational benchmarking revealed that the af2cycler improved TM-scores and reduced PAE for Chroma-generated drafts across a range of protein sizes, without large alterations to the original design. The af2cycler-refined designs folded well and showed their intended shapes when imaged through negative stain electron microscopy.

Furthermore, the af2cycler adds to the observation that protein backbones which were designed utilizing AF2 (either via hallucination (*4*) or the af2cycler) deliver very high designability, even up to lengths of 1000 amino acids. Interestingly, the usage of RFDiffusion partial diffusion as a denoising tool did not increase the designability of Chroma designs as much as the af2cycler did (Supp. Fig. 5).

Moving forward, we anticipate that the af2cycler’s capabilities can be extended by the community towards more complex applications, ranging from multi chain design, binder design or design of multiple protein conformations, building on the open-source integration of af2cycler in the ColabDesign codebase.

## Funding

We acknowledge funding from the European Research Council Advanced Grant (no. 101018465 (to HD)), DFG grants Gottfried Wilhelm Leibniz Program DI1500/3-1 (to HD), TUM Innovation Network Projekt RISE (HD, CF), NIH grant DP5OD026389 (to SO), NSF MCB2032259 (to SO) and Amgen (SO).

## Author contributions

The study was conceived by CF, SO, and HD, with HD and SO providing supervision. Computational studies were carried out by CF, SO, and DS. CF and DS designed proteins. Wet lab experiments were conducted by CF and LF. CF performed nsTEM and analysis. Funding was acquired by HD and SO. The original draft of this manuscript was written by CF, and edited by HD, SO.

## Competing interests

The authors declare no competing interest pertaining to results described in this study.

## Supplementary Figures

**Supplementary Figure 1:**
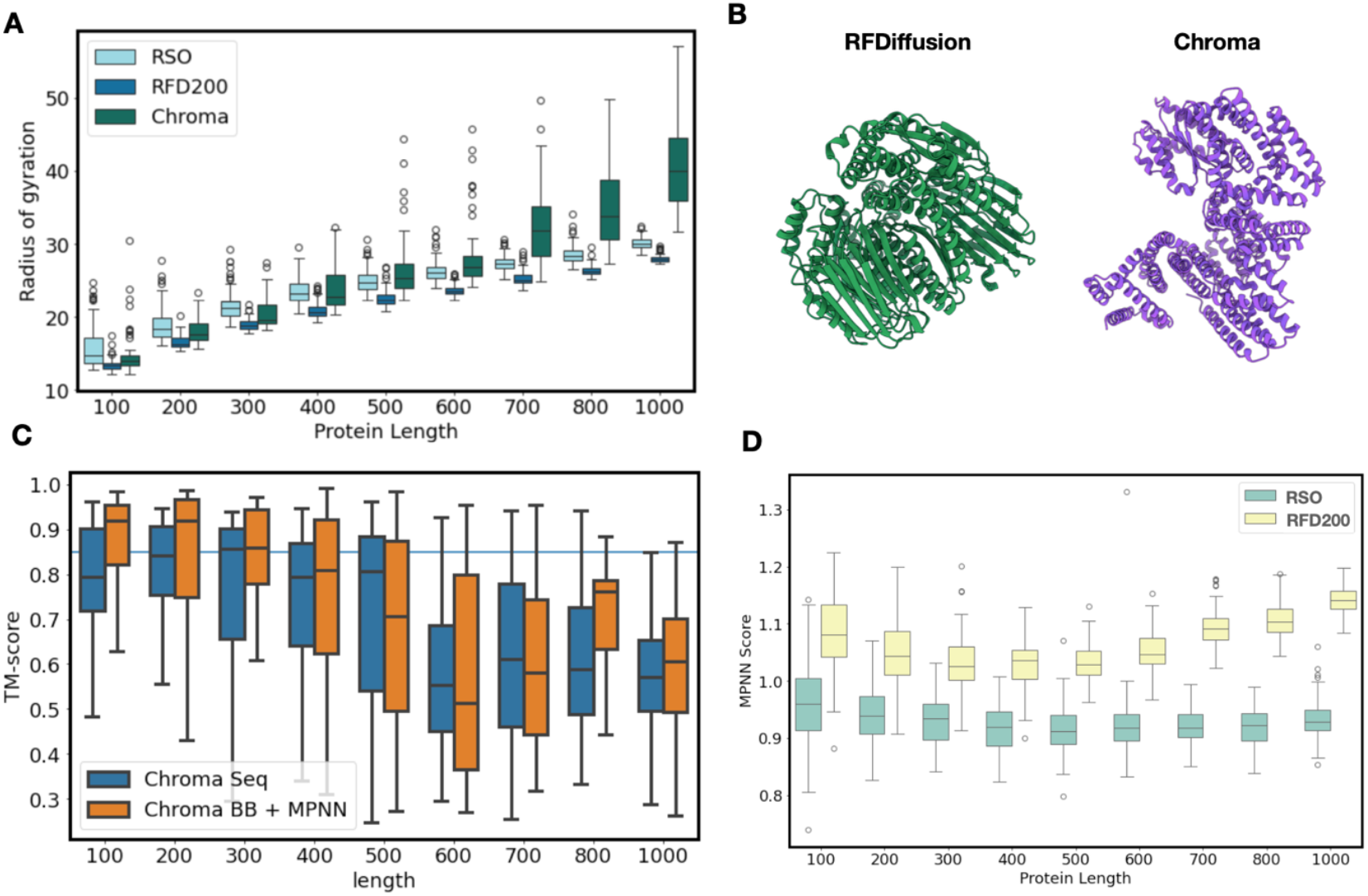
A) Radius of Gyration of protein backbones created either with RSO (cite), RFDiffusion (RFD200) or Chroma. B) Design output from RFDiffusion and Chroma, showing the compactness of RFDiffusion generated designs when compared to Chroma (likely resulting from an inclusion of radius of gyration in the RFDiffusion training) C) Designability TM-score of the ESMFold prediction of unconditional generated samples with Chroma directly (Chroma seq) or with sequences generated from solubleMPNN redesign D) Boxplot showing the ProteinMPNN score of protein backbones designed either with RFDiffusion or RSO (AD)

**Supplementary Figure 2:**
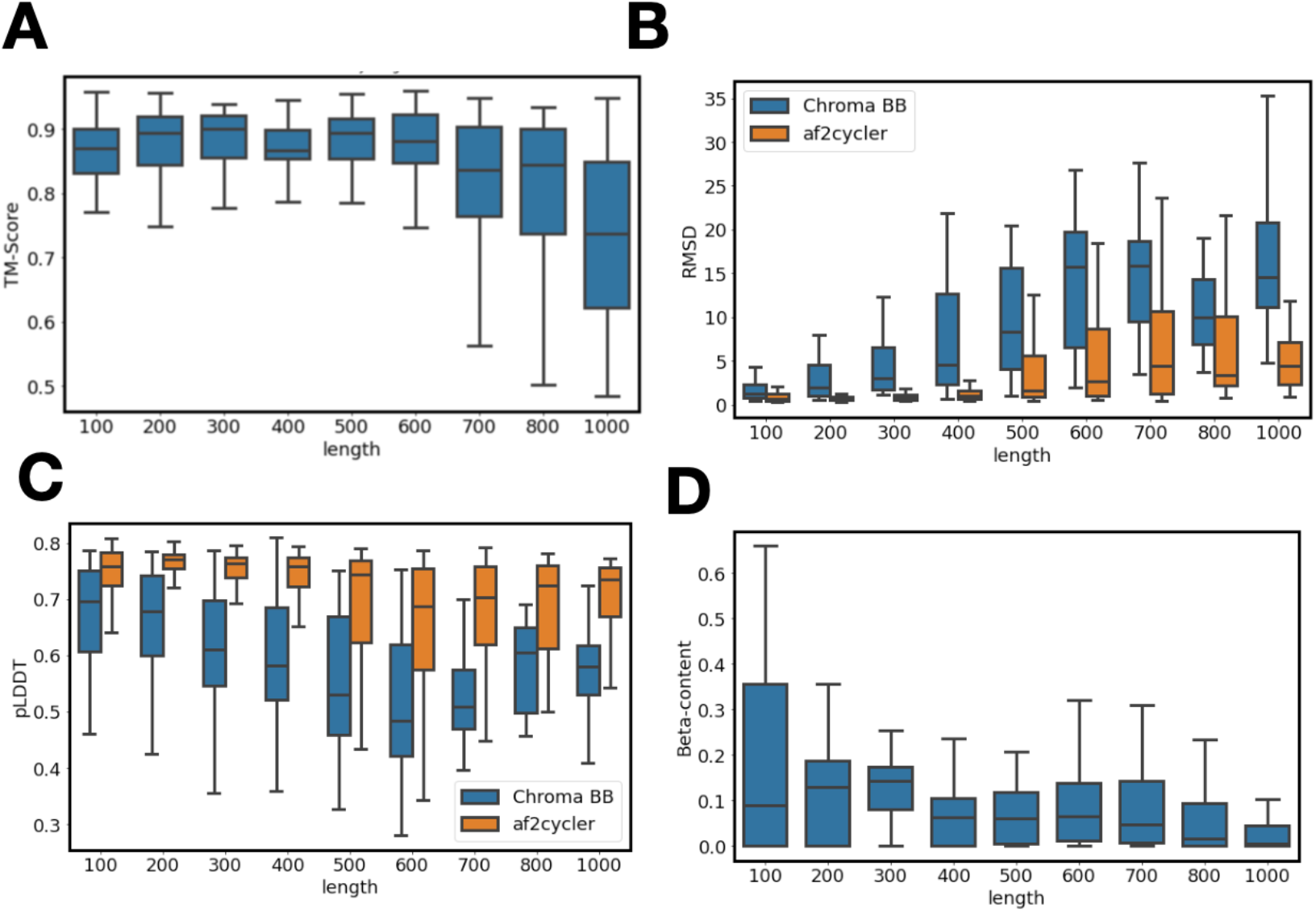
A) Boxplot of the TM-score of Chroma designs before and after af2cycler treatment at certain lengths. B) Designability (RMSD) of Chroma and af2cycler treated backbones C) pLDDT of ESMFold predictions from (B). D) Beta sheet content of chroma generated designs for certain lengths.

**Supplementary Figure 3:**
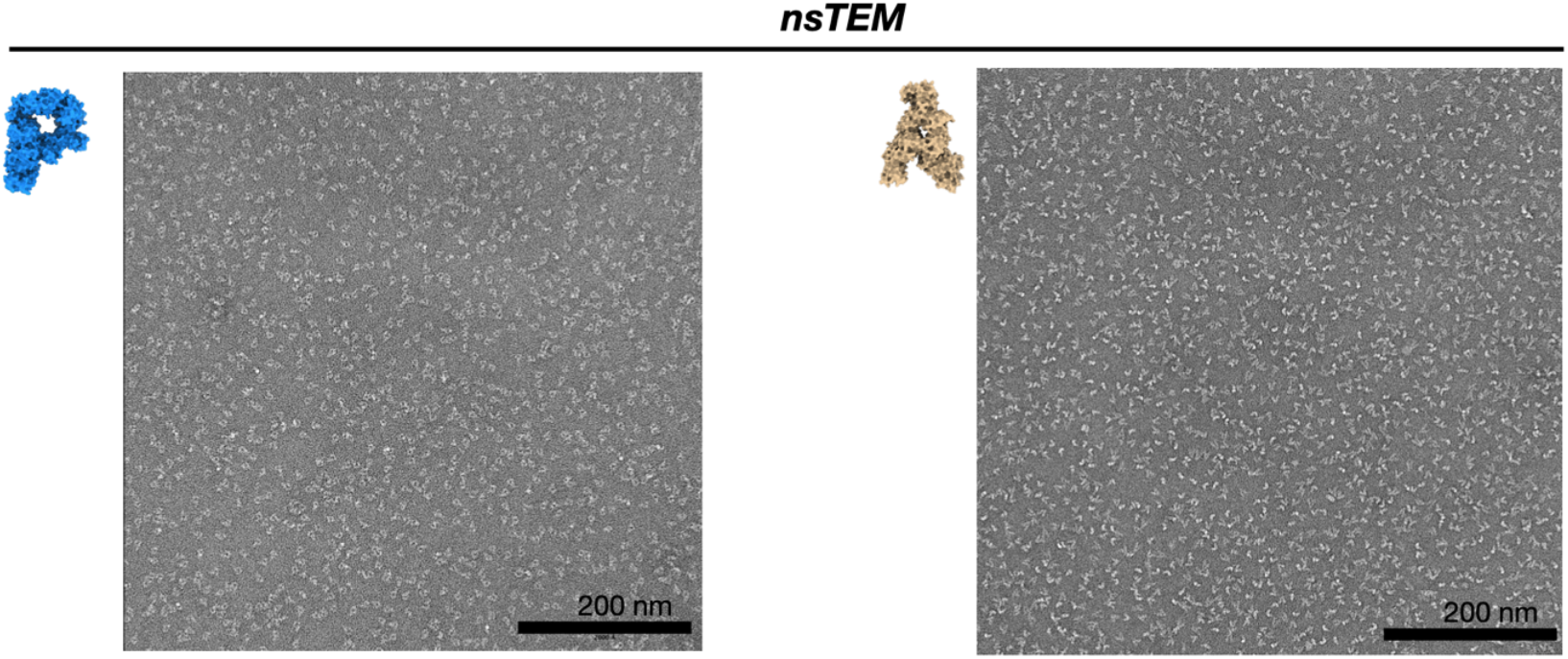
exemplary negative stain TEM micrographs of the ‘P’ and ‘A’ proteins

**Supplementary Figure 4:**
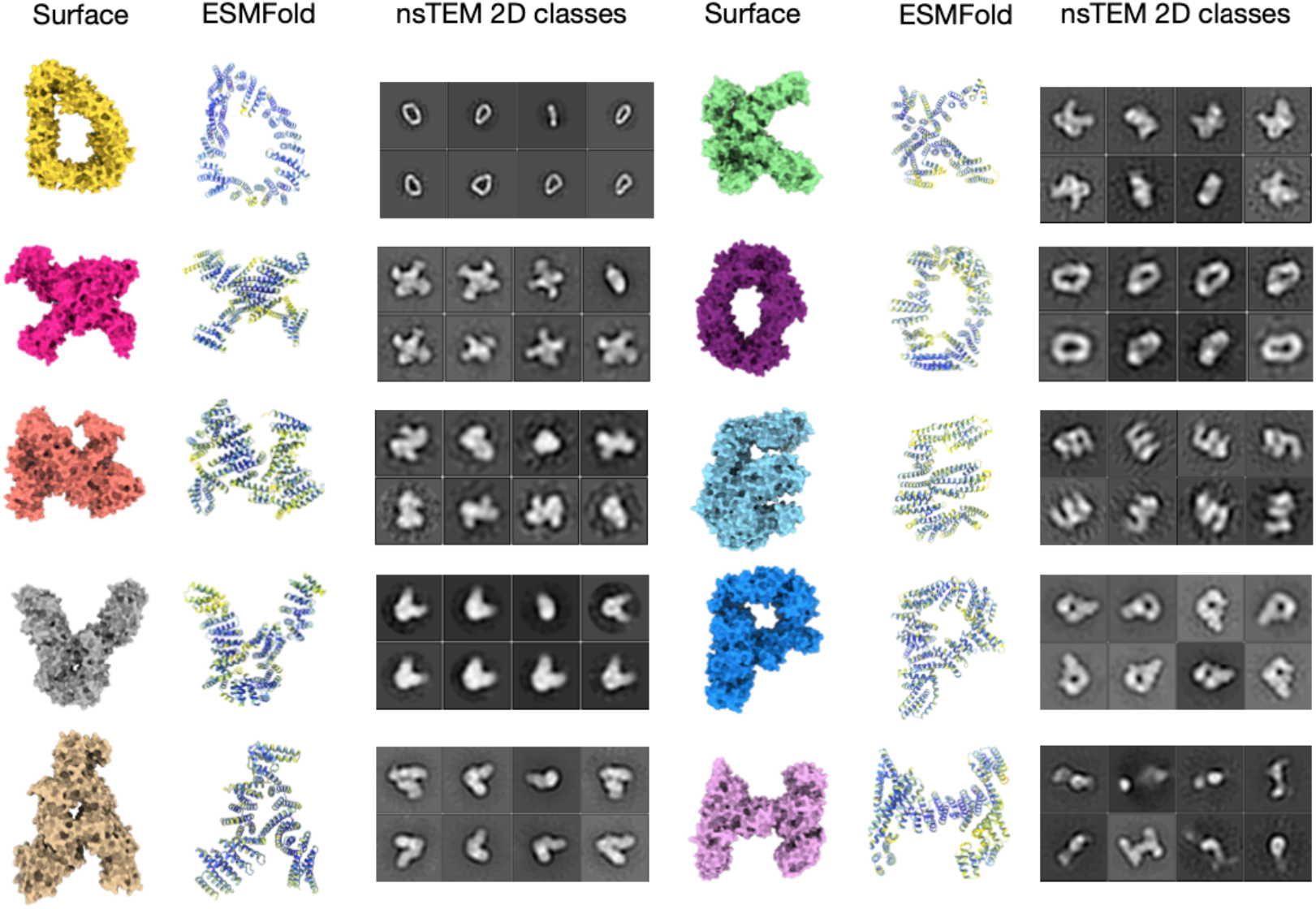
Extended overview of designed letters in surface view, ESMFold predictions colored by pLDDT and extended selection of obtained 2D classes from nsTEM

**Supplementary Figure 5:**
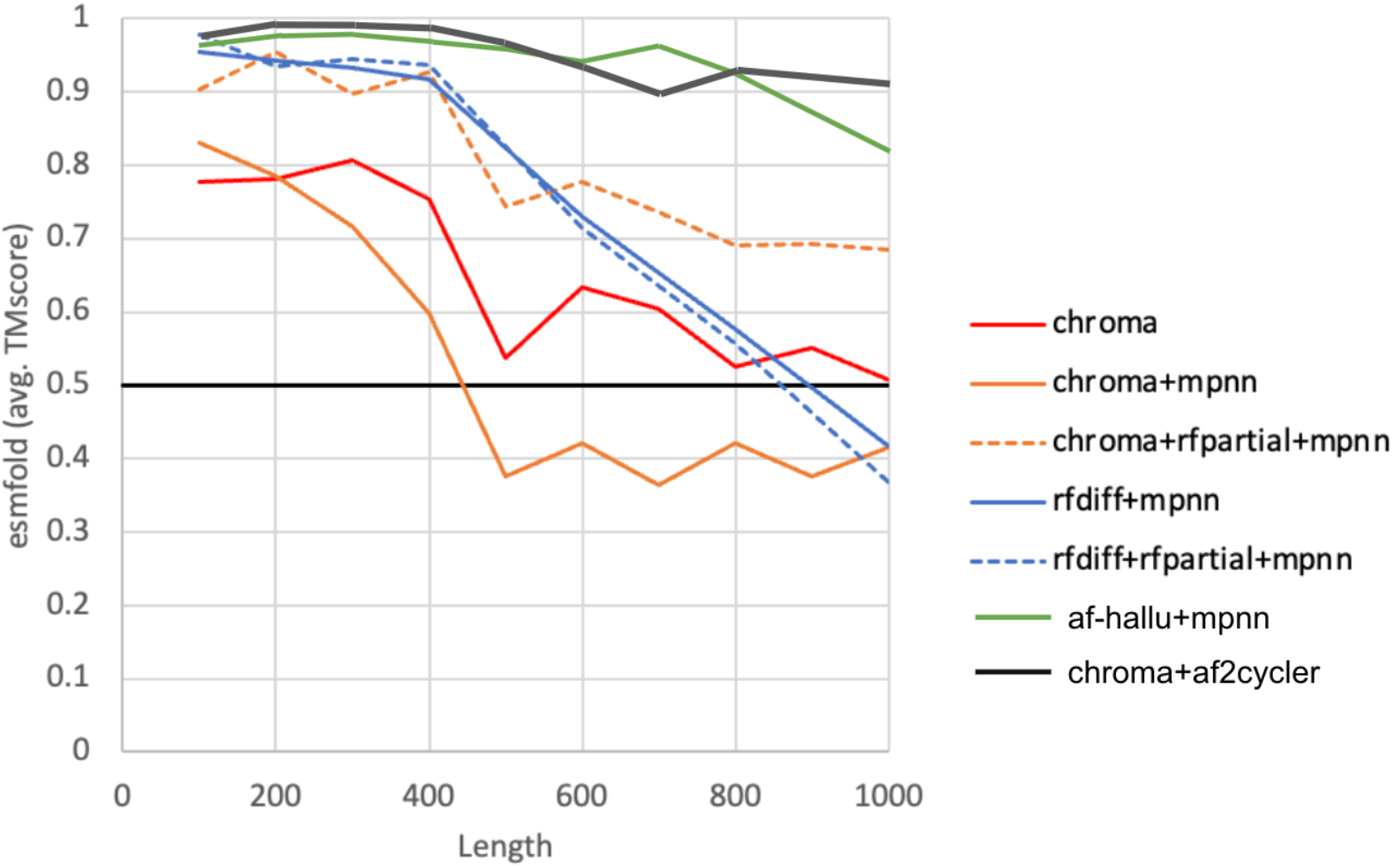
Designability comparison of multiple methods. Unconditional monomeric samples were designed with different methods. Repredictions were performed with ESMFold. **Chroma**: Sequences taken from ChromaDesign **Chroma + MPNN:** Chroma designed backbones were used for ProteinMPNN as input. Eight sequences were generated and repredicted. The sequence with lowest TM Score to the chroma designed backbone was selected. **Chroma + RFDpartial + MPNN**: Chroma designs were subject to partial diffusion (10 steps) using RFDiffusion. Resulting backbones were used as input to ProteinMPNN. **RFDiff + MPNN:** RFDiffusion was used to generate backbones and ProteinMPNN was used to design sequences. **RFDiff + RFDpartial + MPNN**: RFDiffusion was used to generate backbones which were then subject to partial diffusion with RFDiffusion (10 steps) and sequences were generated with ProteinMPNN. **AF-Hallu+MPNN**: Relaxed sequence optimization (*4*) was used to generate backbones and sequences were designed with ProetinMPNN. **Chroma + af2cycler+solMPNN**: Chroma was used to design draft proteins which were used as input into the af2cycler. The resulting backbone was used as input to solubleMPNN

## Materials & Methods

### Chroma design of draft proteins

Chroma draft backbones were design using the available code on GitHub (https://github.com/generatebio/chroma/tree/main) on A100 GPUs. Trajectories were run for either 500 or 1000 steps. Shapes were designed using the “ShapeConditioner” with autoscale_num_residues=1000

### Af2cycler

The af2cycler is implemented using the ColabDesign framework. A notebook with the code to run af2cycling can be found at: https://colab.research.google.com/drive/14ULdrjOmH-XMtGDrikzjDF1FLegZg3-a?usp=sharing

### Designability tests

Resulting backbones after af2cycling were input to a JAX implementation of ProteinMPNN running the solubleMPNN weights. Eight sequences were sampled at a temperature of 0.1. Sequences were predicted using a hugging face implementation of ESMFold (https://huggingface.co/facebook/esmfold_v1) Final TM Scores and RMSDs to the designed backbones were calculated using TM Score. AF3 predictions were performed using the AF3 server.

### Protein Expression and Purification

Amino acid sequences were codon optimized using benchling and divided roughly into three 1000 bp stretches, dependent on position of optimal four base combinations for Golden gate cloning. Linear DNA fragments were ordered from TWIST Bioscience and cloned into the LM670 vector (*6*) replacing the lethal ccdb gene using BSAI restriction enzymes. Cloned vectors were transformed into BL21 (DE3) and plated on a kanamycin containing agar plate. Plates were incubated overnight and inspected for colonies. To verify correct assembly, colony PCRs of the obtained colonies were performed using Primers attaching to the vector right before and behind the insert sites. Positive colonies were grown overnight in 5ml homemade Autoinduction medium (For 1 L: 1x TB media (Carl Roth), 2g lactose, 0.5g glucose), supplemented with Kanamycin) in a 24 well deep well plate covered with breathable film and in another culture of 5 ml LB medium. The next day, bacteria were pelleted and lysed using a homemade lysis buffer 1x B-PER bacterial lysis reagent (Thermo scientific), 20 U/ml DNASE I, 10 mM MgCl2 and 500mM NaCl) for 30 min at RT gently shaking. Cell debris was separated from lysate by centrifugation at 4600g for 15 mins. Supernatant was collected. For Ni-NTA purification 100 ul of ROTI®Garose-His/Ni NTA-Beads (Carl Roth) were applied to 96 well filter plates (ACROPREP, 1um fiber glass) and equilibrated with 5CV Wash buffer (1x PBS, 40 mM Imidazol). Supernatant from lysis was applied and incubated for 3 min. By using a vacuum manifold the supernatant was removed from the beads and subsequently washed with 5 ml of wash buffer. Elution was performed by applying 200 ul of elution buffer (1x PBS, 300 mM Imidazol) and incubating for 5 min. Elutions were then analyzed using SDS page gel electrophoresis to identify clones that produced soluble proteins. For positive clones expressing proteins, cryostock were made by mixing 1ml of the LB culture with 1 ml of 50% glycerin solution, aliquoted and stored at -80 C. The remaining 4 ml of LB culture were pelleted and the plasmid DNA was isolated using the Quiagen MiniPrep kit. Plasmids were finally analyzed using whole plasmid sequencing to verify correct assembly.

A 200 ul aliquot of Cryostocks of sequence verified clones was then incubated in 5 ml LB medium, supplemented with kanamycin, for 5 hours. This preculture was then used to inoculate two 750 ml autoinduction media cultures. Cultures were grown for 16-24 hours at 37 C and then for another 24 hours at 16 C. Resulting cells were pelleted and lysed as described above using the vacuum manifold. Resulting elution fractions were analysed in SDS-page gels and SEC purified using a Äkta running a Superdex Increase 200 10/300.

### Negative stain TEM & image processing

SEC or Ni-NTA purified proteins were diluted to 0.1 mg/ml using PBS. 4 ul of sample were applied to glow discharged Formvar FCF400-CU-50 grids (EMS) and incubated for 30s. Excess sample was blotted and stained with 2% aqueous uranyl formate solution containing 25 mM sodium hydroxide for 30s. Grids were imaged on a FEI Tecnai 120. Typically, a 100 or 200 picture montage was collected at 42-67.000x magnification.

Data processing was done using RELION4. Particles were picked using the gaussian/Laplacian autopicker and subject to multiple (typically 2-4) rounds of 2D classification. Final classes were sorted by number of included particles.

## Notes

### Competing Interest Statement

The authors have declared no competing interest.

## Citations

1. T. Kortemme, De novo protein design—From new structures to programmable functions. Cell 187, 526–544 (2024).

2. N. Ferruz, A. Stein, Computational methods for protein design. Protein Engineering, Design and Selection 37, gzae011 (2024).

3. J. Jumper, R. Evans, A. Pritzel, T. Green, M. Figurnov, O. Ronneberger, K. Tunyasuvunakool, R. Bates, A. Žídek, A. Potapenko, A. Bridgland, C. Meyer, S. A. A. Kohl, A. J. Ballard, A. Cowie, B. Romera-Paredes, S. Nikolov, R. Jain, J. Adler, T. Back, S. Petersen, D. Reiman, E. Clancy, M. Zielinski, M. Steinegger, M. Pacholska, T. Berghammer, S. Bodenstein, D. Silver, O. Vinyals, A. W. Senior, K. Kavukcuoglu, P. Kohli, D. Hassabis, Highly accurate protein structure prediction with AlphaFold. Nature 596, 583–589 (2021).

4. C. Frank, A. Khoshouei, L. Fuβ, D. Schiwietz, D. Putz, L. Weber, Z. Zhao, M. Hattori, S. Feng, Y. de Stigter, S. Ovchinnikov, H. Dietz, Scalable protein design using optimization in a relaxed sequence space. Science 386, 439–445 (2024).

5. C. Goverde, B. Wolf, H. Khakzad, S. Rosset, B. E. Correia, “De novo protein design by inversion of the AlphaFold structure prediction network” (preprint, Bioinformatics, 2022); 10.1101/2022.12.13.520346.

6. B. I. M. Wicky, L. F. Milles, A. Courbet, R. J. Ragotte, J. Dauparas, E. Kinfu, S. Tipps, R. D. Kibler, M. Baek, F. DiMaio, X. Li, L. Carter, A. Kang, H. Nguyen, A. K. Bera, D. Baker, Hallucinating symmetric protein assemblies. Science 378, 56–61 (2022).

7. M. Pacesa, L. Nickel, J. Schmidt, E. Pyatova, C. Schellhaas, L. Kissling, A. Alcaraz-Serna, Y. Cho, K. H. Ghamary, L. Vinué, B. J. Yachnin, A. M. Wollacott, S. Buckley, S. Georgeon, C. A. Goverde, G. N. Hatzopoulos, P. Gönczy, Y. D. Muller, G. Schwank, S. Ovchinnikov, B. E. Correia, BindCraft: one-shot design of functional protein binders. bioRxiv [Preprint] (2024). 10.1101/2024.09.30.615802.

8. O. J. Goudy, A. Nallathambi, T. Kinjo, N. Z. Randolph, B. Kuhlman, In silico evolution of autoinhibitory domains for a PD-L1 antagonist using deep learning models. Proceedings of the National Academy of Sciences 120, e2307371120 (2023).

9. J. Ingraham, M. Baranov, Z. Costello, V. Frappier, A. Ismail, S. Tie, W. Wang, V. Xue, F. Obermeyer, A. Beam, G. Grigoryan, Illuminating protein space with a programmable generative model. bioRxiv [Preprint] (2022). 10.1101/2022.12.01.518682.

10. S. Alamdari, N. Thakkar, R. van den Berg, A. X. Lu, N. Fusi, A. P. Amini, K. K. Yang, Protein generation with evolutionary diffusion: sequence is all you need. bioRxiv [Preprint] (2023). 10.1101/2023.09.11.556673.

11. N. Anand, T. Achim, Protein Structure and Sequence Generation with Equivariant Denoising Diffusion Probabilistic Models. arXiv arXiv:2205.15019 [Preprint] (2022). 10.48550/arXiv.2205.15019.

12. Y. Lin, M. AlQuraishi, Generating Novel, Designable, and Diverse Protein Structures by Equivariantly Diffusing Oriented Residue Clouds. arXiv arXiv:2301.12485 [Preprint] (2023). 10.48550/arXiv.2301.12485.

13. B. L. Trippe, J. Yim, D. Tischer, D. Baker, T. Broderick, R. Barzilay, T. Jaakkola, Diffusion probabilistic modeling of protein backbones in 3D for the motif-scaffolding problem. arXiv arXiv:2206.04119 [Preprint] (2023). 10.48550/arXiv.2206.04119.

14. J. L. Watson, D. Juergens, N. R. Bennett, B. L. Trippe, J. Yim, H. E. Eisenach, W. Ahern, A. J. Borst, R. J. Ragotte, L. F. Milles, B. I. M. Wicky, N. Hanikel, S. J. Pellock, A. Courbet, W. Sheffler, J. Wang, P. Venkatesh, I. Sappington, S. V. Torres, A. Lauko, V. De Bortoli, E. Mathieu, S. Ovchinnikov, R. Barzilay, T. S. Jaakkola, F. DiMaio, M. Baek, D. Baker, De novo design of protein structure and function with RFdiffusion. Nature 620, 1089–1100 (2023).

15. B. Jing, B. Berger, T. Jaakkola, AlphaFold Meets Flow Matching for Generating Protein Ensembles. arXiv arXiv:2402.04845 [Preprint] (2024). 10.48550/arXiv.2402.04845.

16. C. Wang, Y. Qu, Z. Peng, Y. Wang, H. Zhu, D. Chen, L. Cao, Proteus: exploring protein structure generation for enhanced designability and efficiency. bioRxiv [Preprint] (2024). 10.1101/2024.02.10.579791.

17. Z. Lin, H. Akin, R. Rao, B. Hie, Z. Zhu, W. Lu, N. Smetanin, R. Verkuil, O. Kabeli, Y. Shmueli, A. dos Santos Costa, M. Fazel-Zarandi, T. Sercu, S. Candido, A. Rives, Evolutionary-scale prediction of atomic-level protein structure with a language model. Science 379, 1123–1130 (2023).

18. C. A. Goverde, M. Pacesa, L. J. Dornfeld, N. Goldbach, S. Georgeon, S. Rosset, J. Dauparas, C. Schellhaas, S. Kozlov, D. Baker, S. Ovchinnikov, B. E. Correia, Computational design of soluble analogues of integral membrane protein structures. bioRxiv [Preprint] (2023). 10.1101/2023.05.09.540044.

19. J. P. Roney, S. Ovchinnikov, State-of-the-Art Estimation of Protein Model Accuracy Using AlphaFold. Phys. Rev. Lett. 129, 238101 (2022).

20. N. R. Bennett, B. Coventry, I. Goreshnik, B. Huang, A. Allen, D. Vafeados, Y. P. Peng, J. Dauparas, M. Baek, L. Stewart, F. DiMaio, S. De Munck, S. N. Savvides, D. Baker, Improving de novo protein binder design with deep learning. Nat Commun 14, 2625 (2023).

21. H. Schweke, M. Pacesa, T. Levin, C. A. Goverde, P. Kumar, Y. Duhoo, L. J. Dornfeld, B. Dubreuil, S. Georgeon, S. Ovchinnikov, D. N. Woolfson, B. E. Correia, S. Dey, E. D. Levy, An atlas of protein homo-oligomerization across domains of life. Cell 187, 999–1010.e15 (2024).

22. D. Kimanius, L. Dong, G. Sharov, T. Nakane, S. H. W. Scheres, New tools for automated cryo-EM single-particle analysis in RELION-4.0. Biochemical Journal 478, 4169–4185 (2021).

23. J. Abramson, J. Adler, J. Dunger, R. Evans, T. Green, A. Pritzel, O. Ronneberger, L. Willmore, A. J. Ballard, J. Bambrick, S. W. Bodenstein, D. A. Evans, C.-C. Hung, M. O’Neill, D. Reiman, K. Tunyasuvunakool, Z. Wu, A. Žemgulytė, E. Arvaniti, C. Beattie, O. Bertolli, A. Bridgland, A. Cherepanov, M. Congreve, A. I. Cowen-Rivers, A. Cowie, M. Figurnov, F. B. Fuchs, H. Gladman, R. Jain, Y. A. Khan, C. M. R. Low, K. Perlin, A. Potapenko, P. Savy, S. Singh, A. Stecula, A. Thillaisundaram, C. Tong, S. Yakneen, E. D. Zhong, M. Zielinski, A. Žídek, V. Bapst, P. Kohli, M. Jaderberg, D. Hassabis, J. M. Jumper, Accurate structure prediction of biomolecular interactions with AlphaFold 3. Nature 630, 493–500 (2024).

